# Description and whole genome sequencing of *Eikenella exigua* sp. nov., isolated from brain abscess and blood

**DOI:** 10.1101/728162

**Authors:** Kristin A. Stormo, Randi M. Nygaard, Torbjørn S. Bruvold, Grete Dimmen, Paul Christoffer Lindemann, Stina Jordal, Øyvind Kommedal

## Abstract

We herein describe the first novel species within the genus Eikenella since it was established in 1972 by the reclassification of “*Bacteroides corrodens*” to *Eikenella corrodens*. From a poly-microbial brain abscess, we encountered an Eikenella isolate PXX that could not validly be named *E. corrodens*. The isolate grew on blood agar with small, translucent, pitting colonies after 3 days of anaerobic incubation. By reviewing previously collected invasive isolates, we found an additional Eikenella strain EI_02 from a blood culture exhibiting the same properties as PXX. Phylogenetic analyses based on both whole genome and individual house-keeping genes confirmed that the two strains allocate in a phylogenetic cluster separate from *E. corrodens*.

Using specific amplification and sequencing of the Eikenella *nusG*-gene as a method, we further detected the novel Eikenella species in six historic brain-abscesses previously reported to contain *E. corrodens* based on 16S metagenomics. Out of 24 Eikenella whole genome projects available in GenBank, eight cluster together with PXX and EI_02. These isolates were recovered from brain abscess (2), blood (1), bone/soft tissue (3), parotid gland (1) and unknown (1). It remains to be investigated whether the new species can be a cause of endocarditis.

The average nucleotide identity (ANI) value between strain PXX and the *E. corrodens* type strain ATCC 23834 was 92.1% and the corresponding genome-to-genome distance value 47.1%, both supporting the classification of PXX as a novel species. For this species we propose the name *Eikenella exigua*. The *E. exigua* type strain is PXX^T^ (DSM 109756^T^, NCTC 14318^T^).

## Introduction

*Eikenella corrodens* has thus far been the only recognized species within the Eikenella genus (1). The bacterium was first isolated by Eiken in 1958 and originally named *“Bacteroides corrodens*” (2) by its ability to make small depressions into the blood agar surface. Its current name was proposed in 1972 when it was distinguished from *Bacteroides* spp. as a facultative anaerobic bacterium (3). Originally isolated from human sputum, *E. corrodens* is now recognized as a colonizer of the oral mucous membranes, the upper respiratory tract and possibly the gastrointestinal tract (4, 5).

The first case studies of patients infected with *E. corrodens* were reported in the 1960s and 70s (6). In addition to bite wounds and soft tissue infections among injection drug users, *E. corrodens* has been isolated from various clinical sites including brain, liver, lung, bone, joints and the genitourinary tract (4, 5, 7-18). It is often found in poly-microbial purulent infections together with other members of the oropharyngeal microbiota (9).

*Eikenella corrodens* is a member of the HACEK group together with *Haemophilus parainfluenzae*, *Aggregatibacter* spp., *Cardiobacterium* spp. and *Kingella* spp. (1). HACEK-group bacteria are associated with infective endocarditis and appear in approximately 3% of cases. Among these, *E. corrodens* is the least common (19).

From a poly-microbial brain abscess, we isolated a slow growing bacterium (strain PXX) with small translucent colonies that displayed pronounced pitting into the agar. The bacterium produced a caramel odor and grew significantly better under anaerobic and microaerophilic conditions. Partial 16S rRNA gene sequencing (V1-V3) showed > 99% similarity with some *E. corrodens* references, but only 98.0% with the *E. corrodens* type strain NCTC 10596 (NZ_LT906482.1).

The combination of a low partial 16S rRNA gene similarity and growth characteristics untypical for *E. corrodens* suggested a novel and as of yet undescribed Eikenella species. We therefore set out to perform a more thorough genetic and phenotypical characterization of the isolate. As part of this investigation, we also re-evaluated the identity of historical Eikenella isolates from our biobank, as well as previous Eikenella identifications made by 16S metagenomic analysis directly from clinical samples.

## Materials and methods

### Strains and samples

We searched the biobank at the Department of Microbiology at Haukeland University Hospital for Eikenella strains isolated from invasive infections from December 2008 to December 2018. Ten isolates were available; four from blood culture, three from pleural empyema, two from brain abscesses and one from a spondylodiscitis (Table 1). The type strain for Eikenella corrodens was obtained from the American Type Culture Collection, ATCC 23834 (= NCTC 10596). All Eikenella isolates were cultured on blood agar plates, incubated at 35-37°C, in a humidified atmosphere under both strict anaerobic, microaerophilic and CO2-enriched (5%) aerobic conditions.

**Table 1.** Overview of clinical *Eikenella* isolates investigated in this study

| Eikenella isolate | Year of sampling | Clinical specimen/origin (Associated disease/infection) | Co-isolated bacteria |
| --- | --- | --- | --- |
| EI_01 | 2012 | Pus from pulmonary abscess. (Cancer recti, pulmonary metastases). | *Bacteroides thetaiotaomicron*<br>*Escherichia coli* |
| EI_02 | 2010 | Blood culture. (Meningitis based on clinical signs and CSF pleocytosis. Negative CSF-culture. No radiological evidence of brain abscess. TTE<sup>a</sup> unremarkable. Chronic periodontitis.) | None |
| EI_03 | 2014 | Pus from spondylodiscitis. (Aortoduodenal fistula. Related psoas abscess.) | *E. coli*<br>*B. thetaiotaomicron*<br>*Prevotella denticola*<br>*Actinomyces odontolyticus* |
| EI_04 | 2013 | Blood culture. (Suspected UTI. TTE unremarkable. Poor dental hygiene.) | *Pseudomonas aeruginosa* |
| EI_05 | 2013 | Blood culture. (Neutropenic sepsis. Sinus cancer.) | None |
| EI_06 | 2017 | Pus from pulmonary abscess/empyema. (Pulmonary cancer.) | *Streptococcus cristatus* |
| EI_07 | 2016 | Pus from brain abscess. (None.) | *Aggregatibacter aphrophilus*<br>*Streptococcus intermedius*<br>*Fusobacterium nucleatum*<br>*Haemophilus pittmaniae* |
| EI_08 | 2016 | Pus from pleural empyema. (None.) | *Streptococcus anginosus*<br>*Parvimonas micra* |
| EI_09 | 2018 | Blood culture. (Terminal ileitis of unknown origin. Unremarkable TEE<sup>b</sup>.) | None |
| PXX | 2018 | Pus from brain abscess. (Sinusitis) | *A. aphrophilus*<br>*S. intermedius*<br>*P. micra*<br>*Neisseria elongata* |

<sup>a</sup>TTE = transthoracic echocardiograph, <sup>b</sup>TEE = transesophageal echocardiography.

We also included remnant sample-DNA from a recent national Norwegian study on bacterial brain abscesses (20). In that study, six out of 27 brain abscesses with an oral or sinus origin were reported to contain *E. corrodens* by 16S metagenomics whereof only one was successfully cultured. However, bacterial identification in that study was based on the variable areas V1-V2 of the 16S rRNA gene, where the difference between *E. corrodens* and the putative new Eikenella species is small (0% to 0.6%). We therefore sought to reanalyze these six samples (P02, P30, P38, P47, P53 and P67; Table S1) using an alternative gene-target with better discriminative properties.

For the phylogenetic analyses, we also included all Eikenella whole genomes in GenBank. Complete genomes were available for *E. corrodens* type strain (NCTC 10596) and *E. corrodens* KCOM 3110. In addition, 22 Eikenella whole genome projects were available in the GenBank-wgs database (Table 2).

**Table 2.** Overview of *Eikenella* complete genomes and genome projects available from GenBank

| Strain | GenBank accession | Source of isolation | Current name | Proposed name |
| --- | --- | --- | --- | --- |
| 491_NMEN | SAMN03197684 | unknown | *E. corrodens* | *E. corrodens* |
| CC921 | SAMN00103446 | unknown | *E. corrodens* | *E. corrodens* |
| HMSC061C02 | SAMN04495803 | tracheal aspirate | *Eikenella* sp. | *E. corrodens* |
| HMSC071B05 | SAMN04477611 | hip abscess | *Eikenella* sp. | *E. corrodens* |
| HMSC073A11 | SAMN04480449 | hip abscess | *Eikenella* sp. | *E. corrodens* |
| KCOM 1378 | SAMN06947081 | mandibular osteomyelitis | *E. corrodens* | *E. corrodens* |
| KCOM 3110 | NZ_CP034670.1 | subgingival plaque | *E. corrodens* | *Eikenella* sp. (undescribed) |
| NCTC 10596 (T) | NZ_LT906482.1 | sputum | *E. corrodens* | *E. corrodens* |
| NML01-0040 | SAMN04931629 | finger | *E. corrodens* | *E. corrodens* |
| NML01-0328 | SAMN04931630 | hand | *E. corrodens* | *E. corrodens* |
| NML01-A-086 | SAMN04931635 | unknown | *Eikenella* sp. | *E. exigua* |
| NML02-A-017 | SAMN04931636 | blood | *Eikenella* sp. | *Eikenella* sp. (undescribed) |
| NML03-A-027 | SAMN04931637 | submandibular abscess | *Eikenella* sp. | *E. exigua* |
| NML04-0072 | SAMN04931631 | anastomosis | *E. corrodens* | *E. corrodens* |
| NML070372 | SAMN04931638 | bone | *Eikenella* sp. | *E. exigua* |
| NML080894 | SAMN04931639 | breast abscess | *Eikenella* sp. | *E. exigua* |
| NML120348 | SAMN04931640 | brain abscess | *Eikenella* sp. | *E. exigua* |
| NML120819 | SAMN04931632 | finger | *E. corrodens* | *E. corrodens* |
| NML130388 | SAMN04931633 | peritoneal fluid | *E. corrodens* | *E. corrodens* |
| NML130454 | SAMN04931641 | eye | *Eikenella* sp. | *Eikenella* sp. (undescribed) |
| NML140207 | SAMN04931634 | pelvic abscess | *E. corrodens* | *E. corrodens* |
| NML96-A-049 | SAMN04931642 | blood | *Eikenella* sp. | *E. exigua* |
| NML97-A-109 | SAMN04931643 | brain abscess | *Eikenella* sp. | *E. exigua* |
| NML99-0057 | SAMN04931644 | parotid | *Eikenella* sp. | *E. exigua* |

### Antimicrobial susceptibility testing

Susceptibility testing was performed at the EUCAST Development Laboratory for bacteria in Växjö, Sweden.

### MALDI-TOF MS and phenotypic characterization

Matrix-assisted laser desorption ionization–time of flight mass spectrometry (MALDI-TOF MS) was performed using a Microflex mass spectrometer (Bruker Daltonics, Bremen, Germany) with the FlexControl software (version 3.0) and the Biotyper database (version 3.1).

Oxidase and catalase tests were performed according to Clinical Microbiology Procedures Handbook. Other biochemical results were obtained using Vitek 2 with the NH identification card (bioMérieux, Marcy-l’Étoile, France).

### DNA sequencing, genome assembly and phylogenetic analyses

Amplification and Sanger sequencing of partial *nusG*-gene directly from clinical samples: We designed a novel PCR to amplify the near complete Eikenella *nusG* gene (483/540 bp, 89.4%) directly from human clinical samples using primer sequences Forward-5’-AYTCAACDGGYGTTTCTC-3’ and Reverse-5’-GAGARAATGGCTAAARATTGGTA-3’.

Briefly, 2 µl of sample DNA was added to a reaction mixture of 12.5 µl TB Green Premix Ex Taq II (Tli RNaseH Plus) (TaKaRa, Japan), 1.0 µl of each primer (from 10 µM stock solution) and 8.5 µl PCR-grade water. The PCR was run in a 25 µl reaction tube on a SmartCycler (Cepheid, California, USA) real-time apparatus. The thermal profile included an initial activation step at 95°C for 30 seconds(s) followed by 45 cycles of 95°C/10s (melting), 56°C/15s (annealing) and 72°C/20s (extension). Sanger sequencing was performed using the ABI PRISM 1.1 Big-dye sequencing kit and a 3730 DNA Analyzer (ThermoFisher Scientific, Massachusetts, USA).

### Whole-genome sequencing

Bacterial colonies from all study strains were harvested from blood-agar plates and suspended in 400 µl Bacterial Lysis Buffer in a bead-containing tube (SeptiFast LysKit), and run for 2×45 seconds at speed 6.5 at a MagnaLyser apparatus (all Roche, Mannheim, Germany). DNA was extracted using a MagNaPure Compact instrument (Roche) with the Bacterial_NA protocol and an eluate volume of 50 µl. Whole genome sequencing was performed on the Illumina MiSeq platform using the Nextera XT DNA Library Prep Kit and Miseq Reagent Kit v2 with 2×150 cycles according to the recommendations made by the manufacturer. Genome data were assembled using SPAdes (version 3.13.0).

In order to obtain a complete genome for the proposed type strain PXX, supplementary long read sequencing was performed using the MinION platform, the Rapid Sequencing Kit and a FLO-MIN106 vR9 flow cell (all Oxford Nanopore, Oxford, Great Britain). The Unicycler software (version 0.4.7) (21) was used for hybrid assembly of Illumina and MinION data.

### Phylogenetic analyses based on 16S rRNA and housekeeping genes

All analyses were performed in the Geneious software version 9.1.7 (Biomatters, Auckland, New Zealand). Complete 16S rRNA and *rpoB*-genes were extracted from the whole genome projects and aligned using MUSCLE alignment. For the *nusG*-gene, partial sequences corresponding to the *nusG*-PCR amplicons were extracted and aligned together with the *nusG*-sequences obtained directly from the included brain abscesses. Genetic distances were calculated using the Tamura-Nei model and evaluated by bootstrap analysis with 500 replications. The neighbour-joining method was used for construction of phylogenetic trees.

### Whole-genome SNP-based phylogenetic analysis

Whole genome phylogenies were evaluated using the PathoSurv software (Verditra, Bergen, Norway) that runs a variant of the the CSIPhylogeny principle (22). The CSIPhylogeny is a SNP-based method where unassembled sequencing reads are mapped directly to the reference (in this study the complete genome of *E. corrodens* NCTC 10596). The PathoSurv version also allows for inclusion of complete or draft genomes in the analysis by in-silico re-fragmenting these into overlapping sequences of a settable length (150 bp in this study). Average Nucleotide Identity (ANI) values were calculated using the EzGenome ANI Calculator (http://www.ezbiocloud.net/ezgenome/ani) (23). Genome-to-genome distances (GGD) were calculated using the DSMZ genome-to-genome distance calculator (http://ggdc.dsmz.de/home.php) (24).

## Results

### Culture and phenotypic testing

The PXX strain did not produce visible colonies under any conditions after 2 days. After three days, barely visible colonies were present on the plates incubated in a microaerophilic or anaerobic atmosphere. After 5 days, colonies had reached a size of ≤ 0.5 mm. They were translucent with a caramel odor and displayed pronounced pitting of the agar surface. After five days, there was also visible growth on the aerobically incubated plates. MALDI-TOF MS provided a category B consistency for *E. corrodens* (score 1.9). The strain was catalase negative. It was also oxidase negative after 60 seconds and consequently defined as oxidase negative. After 90 seconds a delayed color change occurred. The strain was identified as *E. corrodens* by VITEK2, with a 95% probability and “very good” confidence. The single contradicting reaction was a negative alanine-phenylalanine-proline arylamidase (Table 3). Microscopy revealed short, slender, gram-negative rods, indistinguishable from the *E. corrodens* type strain (2).

**Table 3.** VITEK 2 NH card results for the putative novel species *Eikenella exigua* strains PXX (T) and EI_02 and *Eikenella corrodens* strains ATCC 23831 (T), EI_6, EI_8 and EI_9.

| Biochemical Test (NH card)* | PXX | EI_02 | ATCC 23831 | EI_6 | EI_8 | EI_9 |
| --- | --- | --- | --- | --- | --- | --- |
| **APPA** | - | - | + | + | + | + |
| **BGALi** | - | + | - | - | - | - |
| **ELLM** | + | + | + | + | + | + |
| **LeuA** | + | + | + | + | + | + |
| **ODC** | + | + | + | + | + | + |
| **ProA** | + | + | + | + | + | + |

\*Only tests with at least one positive isolate included. All other biochemical tests were negative for all isolates, including absence of acid production from sugars. APPA = alanine-phenylalanine-proline arylamidase, BGALi = beta-galactopyranosidase Indoxyl, ELLM = Ellman`s reagent, LeuA = leucine arylamidase, ODC = ornithine decarboxylase, ProA = proline arylamidase.

Strain EI_02 showed phenotypic traits similar to PXX except that colonies were visible already after 24h of anaerobic/microaerophilic incubation and that it failed to grow aerobically even after 7 days of incubation. This isolate obtained a category A consistency for *E. corrodens* in MALDI-TOF MS (score 2.1). Catalase and oxidase results were equal to the results for the PXX strain. EI_02 was also reported as *E. corrodens* by VITEK2, but with a lower probability of 89% and only “good” confidence. The contradicting reactions were a negative alanine-phenylalanine-proline arylamidase and a positive beta-galactopyranosidase Indoxyl (Table 3).

The remaining eight strains from our biobank exhibited growth characteristics as described for *E. corrodens* in the literature. Equivalent to the *E. corrodens* type strain ATCC 23834, they produced greyish colonies under all tested atmospheric conditions after 1-2 days, with various degrees of pitting and a characteristic hypochlorite odor. Three of these strains and the type strain of *E. corrodens* were subjected to further phenotypic testing. All four isolates obtained a category A consistency for Eikenella corrodens in MALDI-TOF MS (score 2.1-2.4). They were weakly oxidase positive (within 50-60 sec.) and catalase negative. All were reported as *E. corrodens* by the VITEK 2 NH card with 99% probability and “excellent” confidence with no contradicting reactions (Table 3).

### Antimicrobial susceptibility testing

Both strains belonging to the putative new species (PXX and EI_02) failed to grow in Mueller-Hinton fastidious broth and barely grew on Mueller-Hinton fastidious plates despite prolonged incubation. Both isolates grew well on supplemented Brucella agar and MIC-values were obtained using Etests (bioMérieux).

The susceptibility pattern for the two strains were comparable. Both were susceptible to ampicillin, ceftriaxone, trimethoprim-sulfamethoxazole and ciprofloxacin, susceptible with increased exposure to penicillin-G, and resistant to gentamicin (Table 4).

**Table 4.** Antibiotic susceptibility pattern for *Eikenella exigua* strains PXX (T) and EI_02.

|  | **PXX**<br>**24h MIC**<br>**µg/mL** | **PXX**<br>**48h MIC**<br>**µg/mL** | **EI_02**<br>**24h MIC**<br>**µg/mL** | **EI_02**<br>**48h MIC**<br>**µg/mL** |
| --- | --- | --- | --- | --- |
| **PCG** | 0,5 | 0,5 | 2 | 1 |
| **AMP** | 1 | 1 | 2 | 1 |
| **CRO** | 0,064 | 0,064 | 0,25 | 0,25 |
| **TSU\*** | 0,125 | 0,125 | 0,032 | 0,064 |
| **CIP** | 0,004 | 0,004 | 0,008 | 0,004 |
| **GEN\*** | 64 | 64 | 32 | 64 |

The bacterium was susceptible with increased exposure to penicillin-G (PCG), susceptible to ampicillin (AMP), ceftriaxone (CRO and ciprofloxacin (CIP). There is insufficient evidence (\*) that the organism is a good target for therapy with trimethoprim-sulfamethoxazole (TSU) and gentamicin (GEN). The results were interpreted using PK-PD (Non-species related) breakpoints (EUCAST Clinical Breakpoint Tables v. 9.0, valid from 2019-01-01). PCG was difficult to read after 24h. Etest on Brucella agar (in-house produced plates), 1.0 McFarland, anaerobic incubation for 24 and 48 hours. EUCAST Development Laboratory (EDL) for bacteria, Växjö, Sweden.

### Phylogenetic analyses

For the phylogenetic trees, the main topology was consistent across all comparisons (Figures 1a-d), except for *E. corrodens* 120819 that clustered outside the *E. corrodens* main branch in the 16S rRNA tree. All trees show that our proposed novel type strain PXX is part of a phylogenetic cluster separate from the *E. corrodens* cluster.

**Figure 1a.**
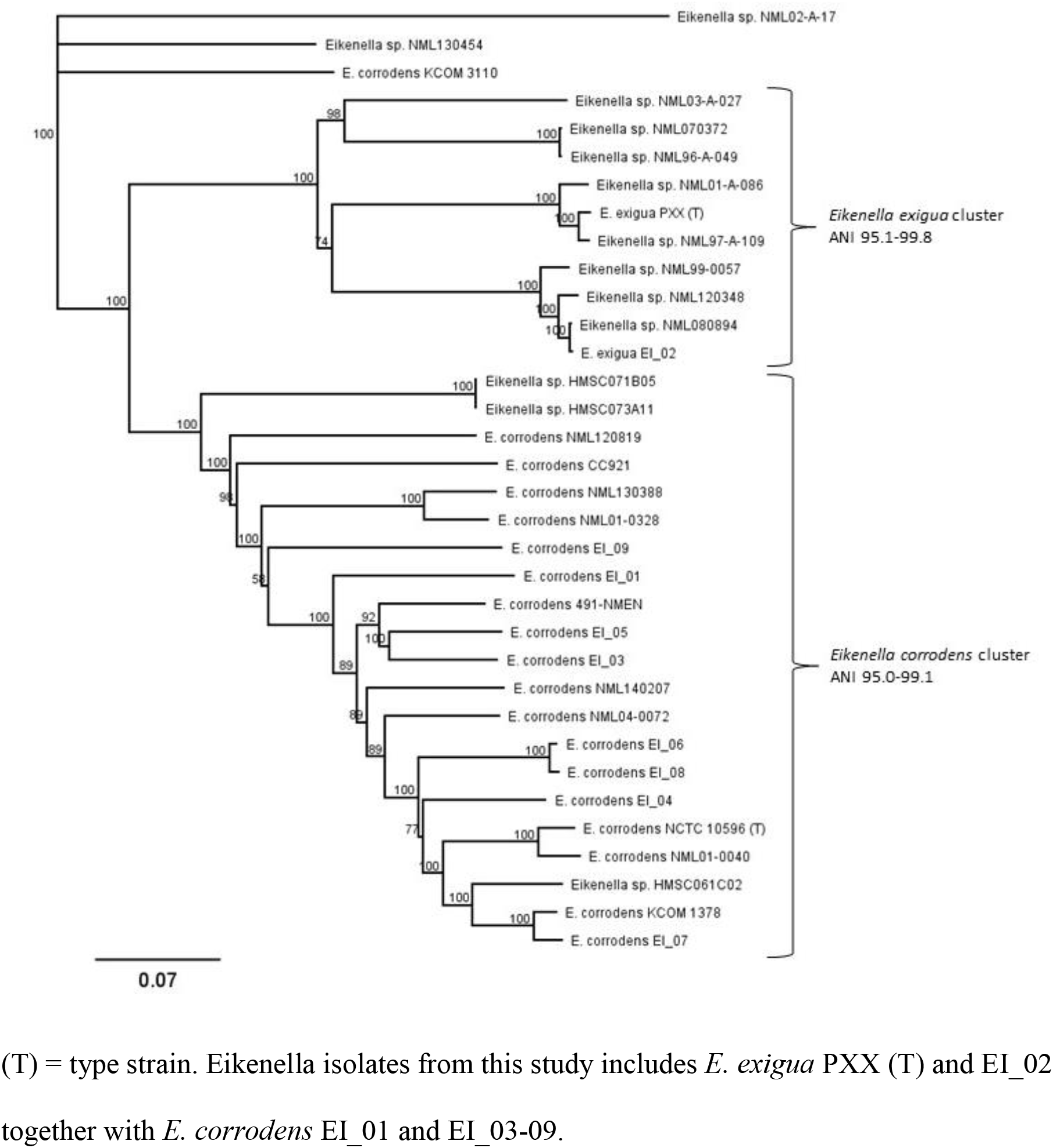
SNP-based whole genome phylogenetic tree

**Figure 1b.**
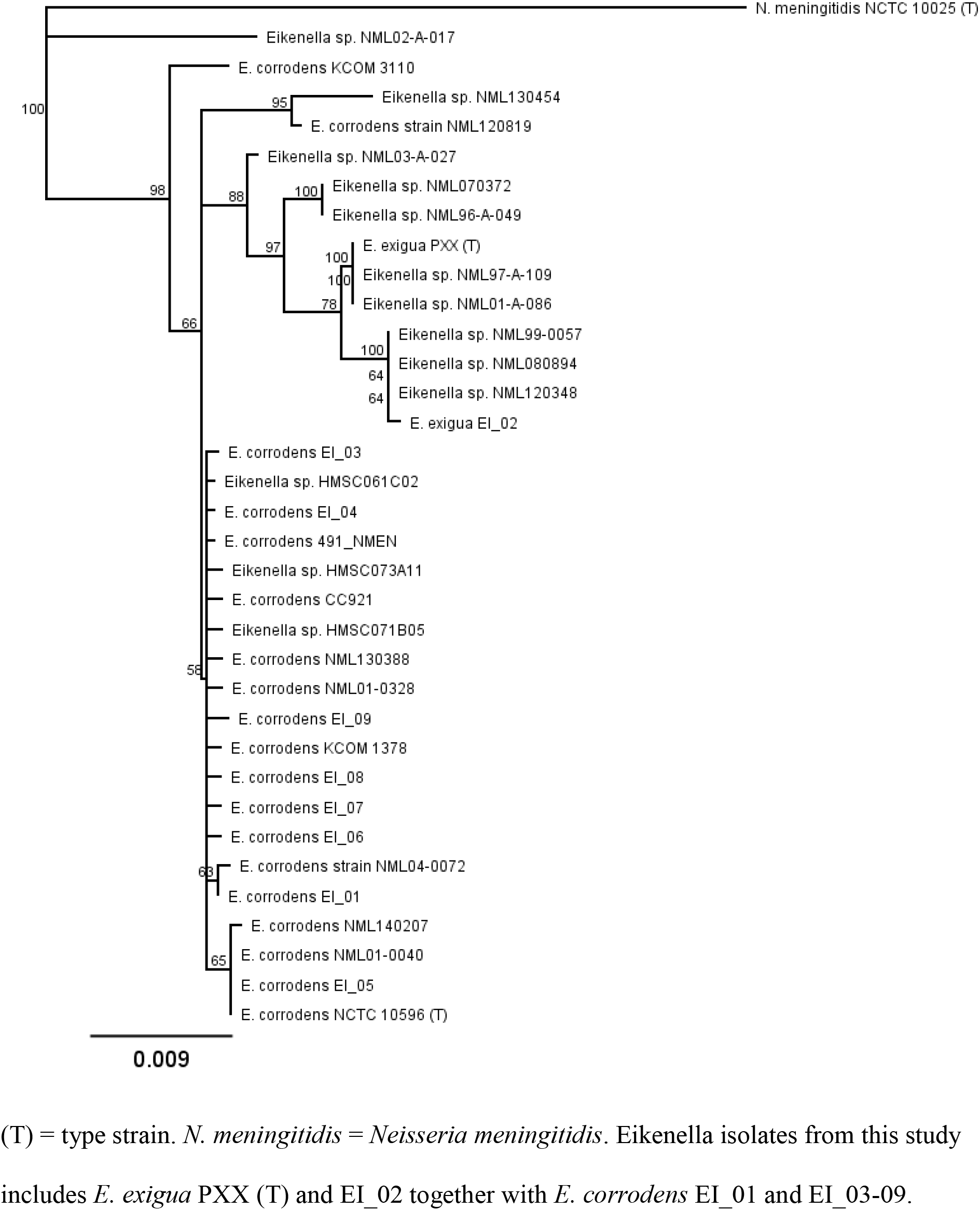
Phylogenetic tree based on the complete 16S rRNA-gene (1499-1500 base pairs)

**Figure 1c.**
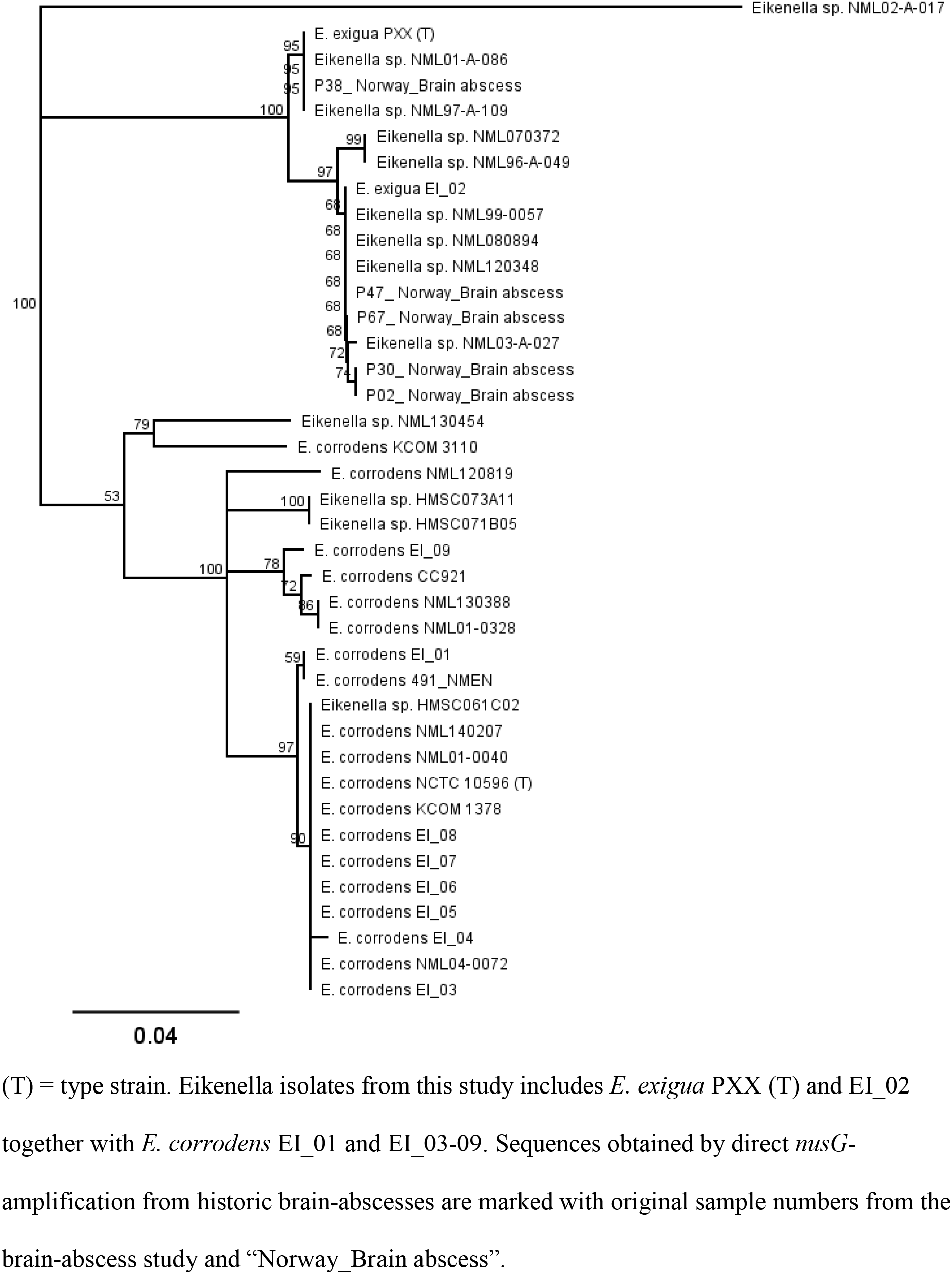
Phylogenetic tree based on partial *nusG*-genes (483 base pairs)

**Figure 1d.**
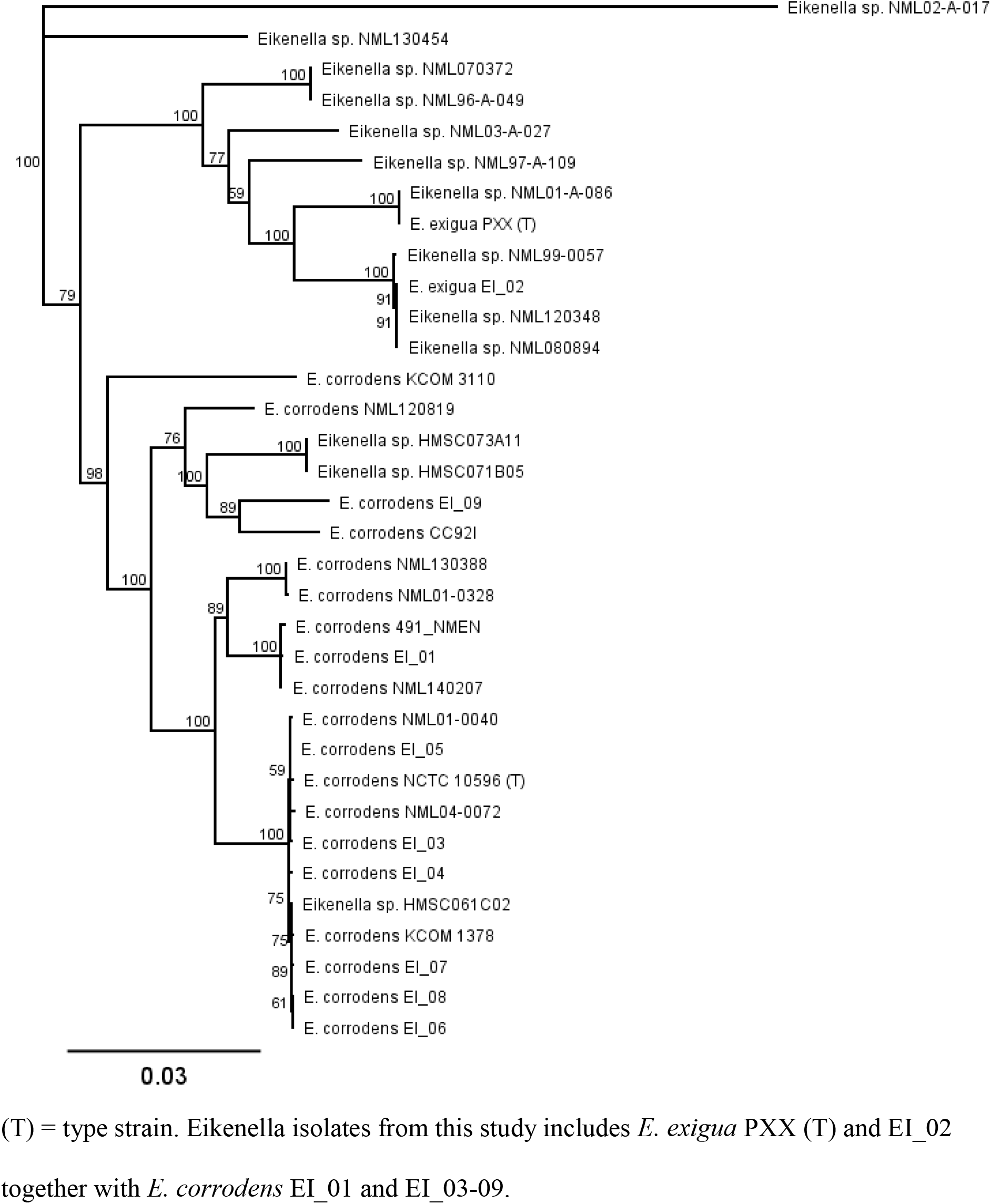
Phylogenetic tree based on the complete *rpoB*-gene (4182 base pairs):

Genome-to-genome distances (GGD) and values for average nucleotide identities (ANI) between representative strains of all putative novel species are provided in Table 5. Both approaches clearly support that the PXX*-*cluster indeed represent a novel species separate from *E. corrodens* for which we propose the name *Eikenella exigua*. The ANI and GGD values between the type strain of *E. corrodens* and the suggested type strain for *E. exigua* were 92.1% and 47.1% respectively. Both the *E. corrodens* cluster and the *E. exigua* cluster were relatively heterogeneous with some intra-cluster ANI-values approaching 95% (Figure 1a). Some GGD values even fell well below 70% in both clusters (range 61.7-92.4% and 60.1-96.1% for the *E. corrodens* and *E. exigua* cluster respectively).

**Table 5:** ANI-values (5a) and GGD values (5b) between representative strains of all separate clusters as indicated in the phylogenetic trees.

a)
|  | PXX (T) | NCTC 10596 (T) | NML02-A-017 | NML 130454 | KCOM 3110 |
| --- | --- | --- | --- | --- | --- |
| PXX (T) |  | 92.1 | 84.7 | 91.3 | 91.2 |
| NCTC 10596 (T) | 92.1 |  | 85.2 | 93.0 | 93.0 |
| NML02-A-017 | 84.7 | 85.2 |  | 85.7 | 85.7 |
| NML 130454 | 91.3 | 93.0 | 85.7 |  | 93.6 |
| KCOM 3110 | 91.2 | 93.0 | 85.7 | 93.6 |  |

b)
|  | PXX (T) | NCTC 10596 (T) | NML02-A-017 | NML 130454 | KCOM 3110 |
| --- | --- | --- | --- | --- | --- |
| PXX (T) |  | 47.1 | 29.5 | 44.2 | 44.9 |
| NCTC 10596 (T) | 47.1 |  | 29.7 | 51.4 | 51.7 |
| NML02-A-017 | 29.5 | 29.7 |  | 30.6 | 42.8 |
| NML 130454 | 44.2 | 51.4 | 30.6 |  | 56.4 |
| KCOM 3110 | 44.9 | 51.7 | 42.8 | 56.4 |  |

Interestingly, our investigations also clearly indicate the existence of three more undescribed Eikenella species among the GenBank whole genomes (Figures 1a-1d and S1, and Tables 6a and 6b), represented by one strain each (NML130454, NML02-A-017 and KCOM 3110 respectively). Table 2 summarizes all GenBank whole-genome sequencing projects investigated in this study, including their origin or associated infection whenever this information was available, their original name in GenBank and their proposed taxonomic classification based on the findings in this study.

### *nusG*-gene sequencing directly from clinical samples

All six brain-abscess reported to contain *E. corrodens* yielded a positive *nusG*-PCR. Sanger sequencing produced unambigous sequences from five samples that were included in the *nusG*-phylogenetic analysis (Figure 1c). All five were representative of *E. exigua*. The sixth electropherogram (from sample P53) contained multiple ambiguous peaks. It was analyzed in the RipSeq mixed software (Pathogenomix, California, USA) and found to represent a mixture of the *nusG*-gene for *E. corrodens* and *E. exigua* respectively.

### Genome properties

Illumina sequencing mean coverage for *E. exigua* strain PXX was 92x, further supplemented with MinION long read sequencing. The complete circular chromosome has a size of 1953249 base pairs and a GC content of 55.5%. It encodes an estimated 2018 genes, including four identical copies of the 16S rRNA gene.

Strain PXX also contains a single 39819 base pair plasmid encoding 19 genes. Except from genes related to the conjugation system and a toxin-antitoxin system, only hypothetical proteins were found on the plasmid. The plasmid was not found in *E. exigua* strain EI_02, but a homologous gen-segment was detected in two GenBank whole-genome projects, one in the *E. exigua* cluster (NML01-A-08, 98,9% homology) and the other probably representing an undescribed Eikenella species (NML130454, 94,8% homology). The plasmid was not found in any of the study *E. corrodens* isolates nor in any *E. corrodens* GenBank whole genome projects.

The GenBank accession numbers for the complete whole-genome sequence of the novel type strain for Eikenella exigua are CP038018 (chromosome) and CP038019 (plasmid). The GenBank accession number for the for the full-length 16S rRNA gene sequence is MK541038.

## Discussion

DNA-DNA hybridization (DDH) experiments have long been the gold standard for interspecies demarcation. More recently, with the introduction of novel sequencing technologies, in silico methods for calculating genomic distances have been accepted as alternatives to DDH.

In this study, we used ANI and GGD to assess relatedness among ten Eikenella isolates and 24 Eikenella whole-genome datasets. The recommended boundaries for a species is 95-96% for ANI and 70% for GGD respectively (23, 24).

Phylogenetic analyses based on bacterial whole genomes and three individual house-keeping genes all showed the existence of an Eikenella cluster separate from the *E. corrodens* cluster. Both ANI and GGD calculations strongly supported that this cluster represents a novel *Eikenella* species (Table 5). This was further supported by different growth characteristics.

The significance of the observed heterogeneity within both the *E. corrodens* and the novel *E. exigua* cluster remains uncertain. It may indicate the existence of separate subspecies within both species but genomic and phenotypic analyses of more isolates are needed to decide upon this matter. However, it should be emphasized that the observed intra-cluster ANI-values are within the recommended boundaries for a species and that both the ANI-values and GGD-values are similar to those reported in a recent phylogenetic study on *Aggregatibacter* spp. (25).

*Eikenella exigua* seems to be associated with CNS-infections. The type-strain was isolated from a poly-microbial brain abscess and strain E_02 was isolated from a blood culture from a patient with suspected bacterial meningitis. *Eikenella exigua* was further identified from the DNA-eluate of six historical brain-abscesses using *nusG*-gene sequencing. Two of the GenBank whole genome projects in the *E. exigua* cluster (NML97-A-109 and 120348) were also brain abscess isolates. In addition, strains within the *E. exigua* cluster have been isolated by others from blood, bone, a breast abscess, a submandibular abscess and the parotid gland. In a recent study we observed striking similarities between the bacterial flora of pleural empyema and that of brain abscesses (26). Two out of 27 primary pleural empyema were found to include *Eikenella* species. These two samples were re-analyzed using the *nusG*-sequencing approach. One was confirmed to contain *E. corrodens* whereas the other contained a mixture of *E. corrodens* and *E. exigua*.

Information about eventual co-isolated bacteria was available for the two *E. exigua* isolates from our hospital, the brain abscess study (20) and the pleural empyema study (26). Except from the blood-culture isolate EI_02, all identifications of *E. exigua* were in poly-microbial infections (Table 1 and Table S1).

In a recent 20-year retrospective study on endocarditis from our region comprising 706 cases, we did not encounter a single Eikenella-endocarditis (27). It therefore remains to be determined whether *E. exigua* has the ability to cause endocarditis like *E. corrodens*. We encourage laboratories with access to *Eikenella* isolates from infective endocarditis to re-evaluate their species designation with regards to our findings. A fruitful approach for a correct identification might be sequencing of the partial *nusG-*gene as described in this publication, either from cultured isolates or directly from extracted heart valves.

### Description of *Eikenella exigua* sp. nov

*Eikenella exigua* (ɛkˈsigu:ɑ); latin adjective nominative feminine singular for small, meager or sparse referring to the growth characteristics of the bacterium.

*Eikenella exigua* is a facultatively anaerobic, short and slender Gram-negative rod. Colonies are visible on blood agar after 1-3 days of incubation in a microaerophilic or anaerobic atmosphere. It grows poorly (after 5 days) or not at all under aerobic conditions. Colonies are small (≤0.5 mm) and translucent with a caramel odor, and display pronounced pitting of the agar surface. It is catalase and oxidase negative. A negative alanine-phenylalanine-proline arylamidase test can be used to discriminate it biochemically from *E. corrodens*. It can also be distinguished from *E. corrodens* based on partial 16S rRNA sequencing including e.g. the variable areas V1-V3 or V3-V4. The genome size is 1.95 MB, with a GC content of 55.5%.

*Eikenella exigua* has been detected in samples from several invasive infections including brain abscesses, blood, bone and pleural empyema. It has also been isolated from a submandibular abscess and the parotid gland and is probably a commensal of the human oropharyngeal microbiota.

The type strain is *Eikenella exigua* PXX^T^ (DSM 109756^T^, NCTC 14318^T^) and was isolated from a poly-microbial brain abscess at Haukeland University Hospital, Bergen, Norway in 2018.

## Acknowledgements

We would like to thank Erika Matuschek at the EUCAST Development Laboratory (EDL) for bacteria, Central Hospital, Växjö, Sweden for performing the susceptibility testing.

We would also like to thank Dr. Sindre Horn for his excellent assistance in the process of choosing a correct and proper Latin name for the novel species.

